# Association of Metabolic and DNA-repair gene polymorphisms with Longevity: a role for *GSTT1, GSTP* and *XPC* genes

**DOI:** 10.1101/2020.07.28.225433

**Authors:** Manuel Scarfò, Chiara Sciandra, Alfredo Santovito

## Abstract

Aging and longevity are complex processes controlled at different levels, including genetic level. We evaluated the association of seven drug and DNA-repair gene polymorphisms with longevity in an Italian cohort. A sample of 756 subjects aged 18-98 was genotyped for *CYP1A1 exon 7 A>G, GSTT1 null, GSTM1 null, GSTP A>G, XRCC1* exon 6 C>T, XRCC1 exon 9 A> G and XPC exon 15 A>C gene polymorphisms. The association between the analyzed gene polymorphisms and longevity was evaluated by dividing the sample into three age groups: 10-50, 51-85, and 86-98.

We observed a significant decrease in the frequency of the *GSTT1* null, *GSTP* G and *XPC* C alleles in the oldest group with respect to the youngest one and with respect to 51-85 age group. We obtained the same results also subdividing the sample into 1-85 and 86-98 age groups. The general linear model analyses confirmed a significant decreasing trend of the above mentioned alleles with age. We hypothesized that these minor alleles, being important in the sensitivity against the development of different types of cancer, may reflect a reduced life-expectancy in carrier subjects and may explain their significantly lower frequency observed among subjects belonging to oldest age group.

## 1. Introduction

Longevity and aging are complex processes that are controlled by both environmental (principally lifestyle, including dietary habits) and genetic factors. These last are referred to those gene polymorphisms that play a significant role in homeostasis processes (Budovsky et al., 2013; Santovito, et al., 2019). A favorable environment and healthy lifestyle allow genetically predisposed subjects to increase their longevity and, in general, to reach a satisfactory aging status (Minciullo et al., 2016). Long-lived individuals seem to be able to avoid age-related diseases. Therefore, the evaluation of genes modulating susceptibility to age-related diseases in global population could be a useful target for human longevity studies.

Most of the published papers about genetic basis of the longevity were principally focused on those gene polymorphisms related to susceptibility to cardiovascular diseases, such as *FTO, ACE* and *ApoE* (Santovito et al., 2019; Kuo et al., 2020), while the relationships between drug and DNA-repair gene polymorphisms and longevity was poorly studied through populations studies.

Drug metabolizing enzymes (DMEs) play a key role in protecting the human body. DMEs contribute to metabolism, biotransformation, elimination and/or detoxification of endogenous and xenobiotic compounds to which humans are exposed, most of them can cause several injuries and, since many of them are lipophilic, they remain unionized in the organism (Crocco et al. 2019). For these reasons, different tissues and organs are provided with various DMEs including phase I, phase II metabolizing enzymes and phase III transporters (Xu et al. 2015). In particular, phase I and phase II enzymes permit the biotransformation of a lipid-soluble xenobiotic or endobiotic compound into a more hydrophilic metabolite through functionalization reactions (Phase I) and/or conjugation (Phase II) reactions (Wauthier et al. 2007).

Phase I DMEs mainly consist of the cytochrome P450 (CYP) superfamily of microsomal enzymes (Xu et al. 2015), that catalyze various reactions, but oxidation is the primary (Almazroo et al. 2017).

Phase II DMEs comprise many superfamily of enzymes including glutathione S-transferases (GST). Phase II DMEs increases, through conjugation reactions, the hydrophilicity of xenobiotic or endobiotic compounds (Xu et al. 2015). At the end of the biotransformation process, phase III transporters help in transferring the more hydrophilic products outside the cells (Almazroo et al. 2017).

In order to prevent the potentially mutagenic consequences of DNA modifications following the exposure to environmental xenobiotics, cells have evolved different mechanisms of DNA repair, depending on the specific type of DNA damage. DNA damage is continuously repaired through activation of different mechanisms that involve polymorphic enzymes. These mechanisms include Base Excision Repair (BER) and Nucleotide Excision Repair (NER) that correct non-bulky damage and lesions that distort the DNA double helical structure, respectively (Cleaver et al., 2009; Collins and Azqueta, 2012). DNA-repair genes, being responsible for preventing the integrity of the genome, may be considered as longevity-associated genes, also considering the fact that a reduced repair capacity has been reported to be associated with cancer development (Langie et al., 2015).

Given these premises, in the present study we decided to evaluate the frequencies of seven polymorphic metabolic genes in a sample of the Italian population and the variation of these frequencies in different age groups. We considered the following metabolic genes, that represent the most studied cancer-associated gene polymorphisms: Cytochrome P450 1A1 (*CYP1A1*), Glutathione S-transferases (*GST*) *T1, GSTM1, GSTP*, X-ray repair cross-complementing group 1 (*XRCC1*) and Xeroderma pigmentosum complementation group C (*XPC*) genes.

The *CYP1A1* gene, which encodes a member of the Phase I *CYP* superfamily of enzymes, is primarily expressed in extrahepatic tissues and plays a central role in the metabolism of polycyclic aromatic hydrocarbons (Androutsopoulos et al. 2009). Its gene product can catalyze different reactions, for example it participates in the conversion of environmental carcinogens into their final DNA-binding carcinogenic form (Sen and Stark, 2019). Analysis on the human gene *CYP1A1* sequence demonstrated that the gene is highly polymorphic (Sen and Stark, 2019). In this study we considered *CYP1A1* exon 7 polymorphisms due to a single nucleotide polymorphism in which adenine is replaced by guanine (Wei and Hu 2015). Many studies reported that the resulting amino acid substitution *CYP1A1* Ile(462)Val is associated with a higher risk of various types of cancer (Lòpez-Cima et al. 2012; Wei and Hu 2015).

*GST* genes are widely distributed in nature and codify for one of the major groups of phase II detoxifying enzymes, evolved to protect organisms against toxic substances that come from food and environment (Nebert and Vasiliou, 2007). Human *GSTs* embody members belonging to seven classes, including Mu (*GSTM*), Pi (*GSTP*), and Theta (*GSTT*). *GST* genes are organized in chromosomal clusters and most of them are polymorphic. These polymorphisms also include the complete absence of *GSTM1* and *GSTT1* (Nebert and Vasiliou, 2007), due to the deletion of a segment of DNA, with the subsequent lack of protein synthesis in homozygous individuals. Deletion polymorphism in *GST* genes (non-functional alleles) has been found to be associated with the development of certain types of cancer, such as oral cancer (Saravani et al., 2019).

GSTP1 is the major isoenzyme expressed in human lung tissue. The single nucleotide polymorphism changes an adenine with a guanine in codon 105, resulting in the replacement of an isoleucine with a valine in the enzyme (Lòpez-Cima et al., 2012). Particular attention has been focused on gene coding for these enzymes because polymorphisms are linked with an increase of diseases risk (Poukeramat et al., 2020; Khabaz, 2014).

The *XRCC1* gene encodes the proteins having an important role in the repair of single-strand DNA breaks associated with free oxygen radicals, radiation, ultraviolet, and alkylating agents (Caldecott, 2019). In particular, *XRCC1* gene is involved in the BER pathway, and its two most common single nucleotide polymorphisms (SNPs), Arg194Trp on exon 6 (rs1799782, C>T) and Arg280His on exon 9 (rs25489, A> G) have been associated with bladder cancer (Fang et al., 2013; Li et al., 2013).

Finally, an important key gene involved in the NER pathway is XPC, that encodes a 940 amino-acid protein involved in DNA damage recognition, DNA damage induced cell cycle checkpoint regulation and apoptosis (Fontana et al., 2008). One of the most common polymorphic sites of the *XPC* gene, represented by A>C transition in exon 15 resulting in a lysine-to-glutamine transition at position 939 (Lys939Gln; rs2228001), has been associated with critical events in human bladder cancer carcinogenesis (Chen et al., 2007; Fontana et al., 2008).

## 2. Materials and Methods

### 2.1. Subjects

In this study, we included 756 subjects (aged 10-98 years; 335 males and 421 females) belonging to the population of Northern Italy.

We recruited subjects, who were natives of Northern Italian localities for at least two generations, and voluntarily joined the project. In order to avoid selection bias, subjects were sampled without *a priori* exclusions. All data from each participant, including ancestry, were collected during an interview in an open-ended manner. All subjects received detailed information about the study and gave their informed consent. The possible association of the studied genetic polymorphisms with longevity was evaluated by dividing the sample into three age groups: 10-50, 51-85, and 86-98, representing individuals in reproductive phase, post-reproductive phase, and long-lived individuals, respectively. The research protocol was approved by the local ethics committee and was performed in accordance with the ethical standards laid down in the 2013 Declaration of Helsinki.

### 2.2. DNA Extraction and Genotyping

Heparinized vacutainers were used to collect 5-10 mL of peripheral blood from each subject. Then, the vacutainers were stored at -20°C prior to analysis.

DNA extraction was conducted using salting-out procedure: 0.5 mL of peripheral blood was added to red cell lysis buffer (10 mM Tris pH 7.6; 5 mM MgCl 2 and 10 mM NaCl), gently shake for 30 sec, and centrifuged at 14,000 rpm for 1 min. White blood cells were centrifuged at 14,000 rpm and the pellet was re-suspended in a solution consisting of 340 µL di white cell lysis buffer (10 mM Tris pH 7.6; 10 mM EDTA and 50 mM NaCl), 10 µL of SDS 10% and 30 µL of proteinase K. After incubation at 55°C for 30 min, 200 µL of saturated sodium acetate were added to the solution. Samples were vigorously shaken and centrifuged at 14,000 rpm for 5 min. At the supernatant solution, transferred in a new eppendorf, were added 0.5 mL of isopropanol for DNA precipitation and, successively, after centrifugation at 14.000 rpm for 1 min, 0.5 mL of 70% ethanol were added to remove salt from the pellet. After 30-60 min at room temperature, the pellet was re- suspended in 50 µL of ultrapure distilled water. PCR reactions were performed in a reaction volume of 25 μL, containing 1X reaction buffer, 1.5 mM MgCl2, 5% DMSO, 250 µM dNTPs, 0.5 μM of each primer, and 1 U/sample of Taq DNA polymerase (Fischer Scientific, Rodano (MI), Italy). The PCR cycles were set as follows: 35 cycles, 1 min at 95°C, 1 min at 60°C, 1 min at 72°C, and a final extension step of 10 min at 72 °C. The amplification products were electrophoresed using 2.5% agarose gel and detected by ethidium bromide staining. Primer sequences, melting temperatures, PCR methodologies used, and expected PCR product sizes are reported in Table 1.

**Table 1.**
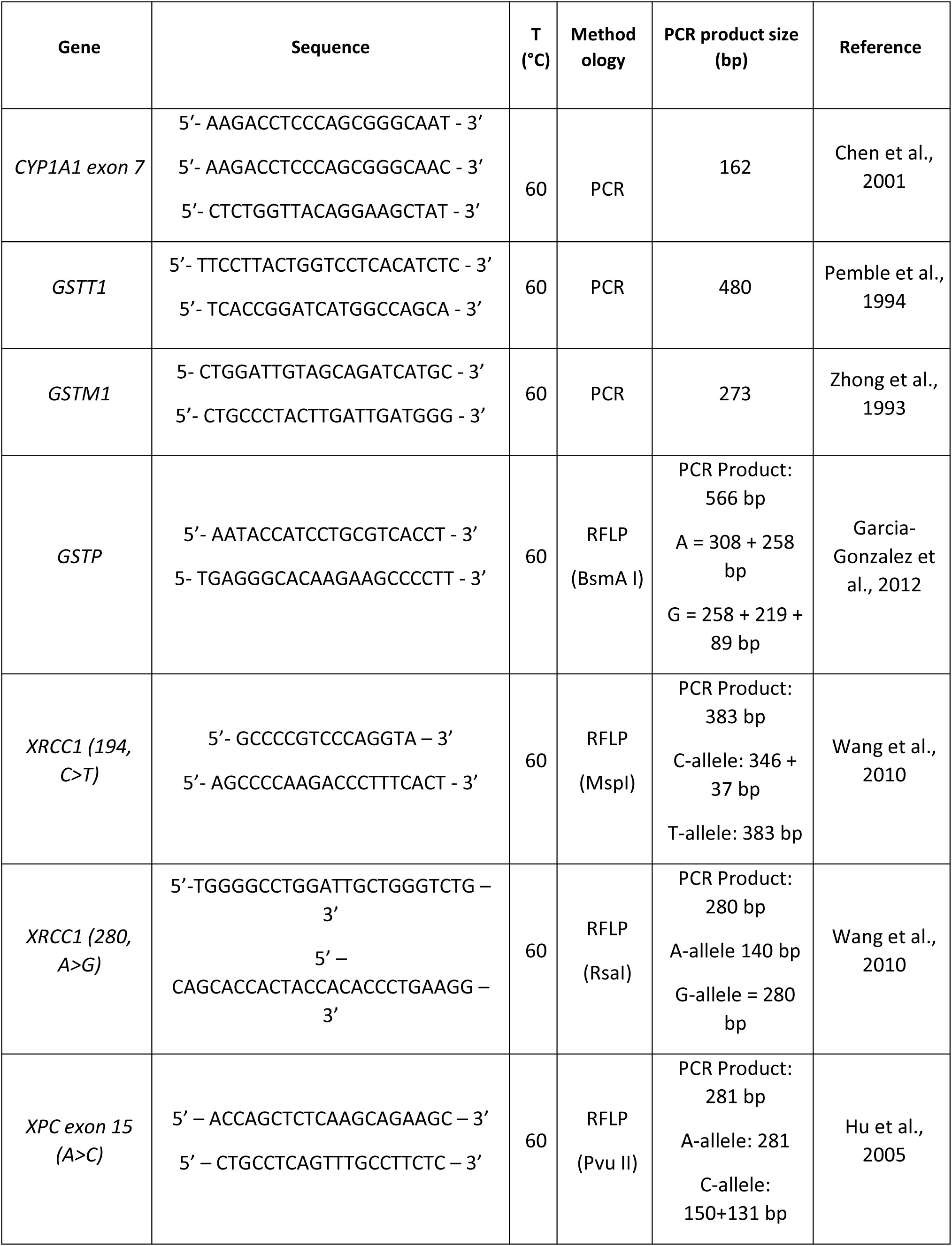
Primers and annealing temperatures for gene polymorphisms analysed in the present study

One hundred random samples were re-analyzed by another investigator in order to verify the accuracy of the genotyping results. The two analyses showed identical results.

### 2.3. Statistical Analysis

All statistical analyses were performed using the SPSS software statistical program (version 25.0, SPSS Inc., Chicago, USA). Pearson’s χ2 test contingency table was used to evaluate both the Hardy–Weinberg equilibrium (HWE) and the statistical differences between two age groups at a time. Comparison of genotype frequencies between age groups was carried out using the Fisher’s exact test. The relationship between age and a specific genotype was analyzed using multivariate general linear model with Bonferroni’s correction. All *P*-values were two-tailed, and the level of statistical significance was set at *P*<0.05 for all tests.

## 3. Results

The general characteristics of the studied sample are reported in Table 2. We recruited 756 subjects, including 335 males (mean age ± SD: 47.499±17.650, age range: 18-94) and 421 females (mean age ± SD: 46.076±21.624, age range: 19-98).

**Table 2.** General characteristics of the studied population.

| **Subjects** | **N (%)** | **Age, Mean ± S.D.**<br>**(range)** |
| --- | --- | --- |
| **Total** | **756** | **55.879±22.676** |
| **Sex** |  |  |
| Males | 335 (44.31) | 47.499±17.650<br>(18-94) |
| Females | 421 (55.69) | 46.076±21.624<br>(19-98) |
| **Age groups** |  |  |
| 18-50 | 538 (71.17) | 35.797±7.799 |
| 51-85 | 151 (19.97) | 66.424±11.566 |
| 86-100 | 67 (8.86) | 89.866±2.639 |

S.D. = Standard Deviation

Allele and genotype frequencies of the analyzed gene polymorphisms are reported in Table 3. All gene polymorphisms were in the HWE.

**Table 3.**
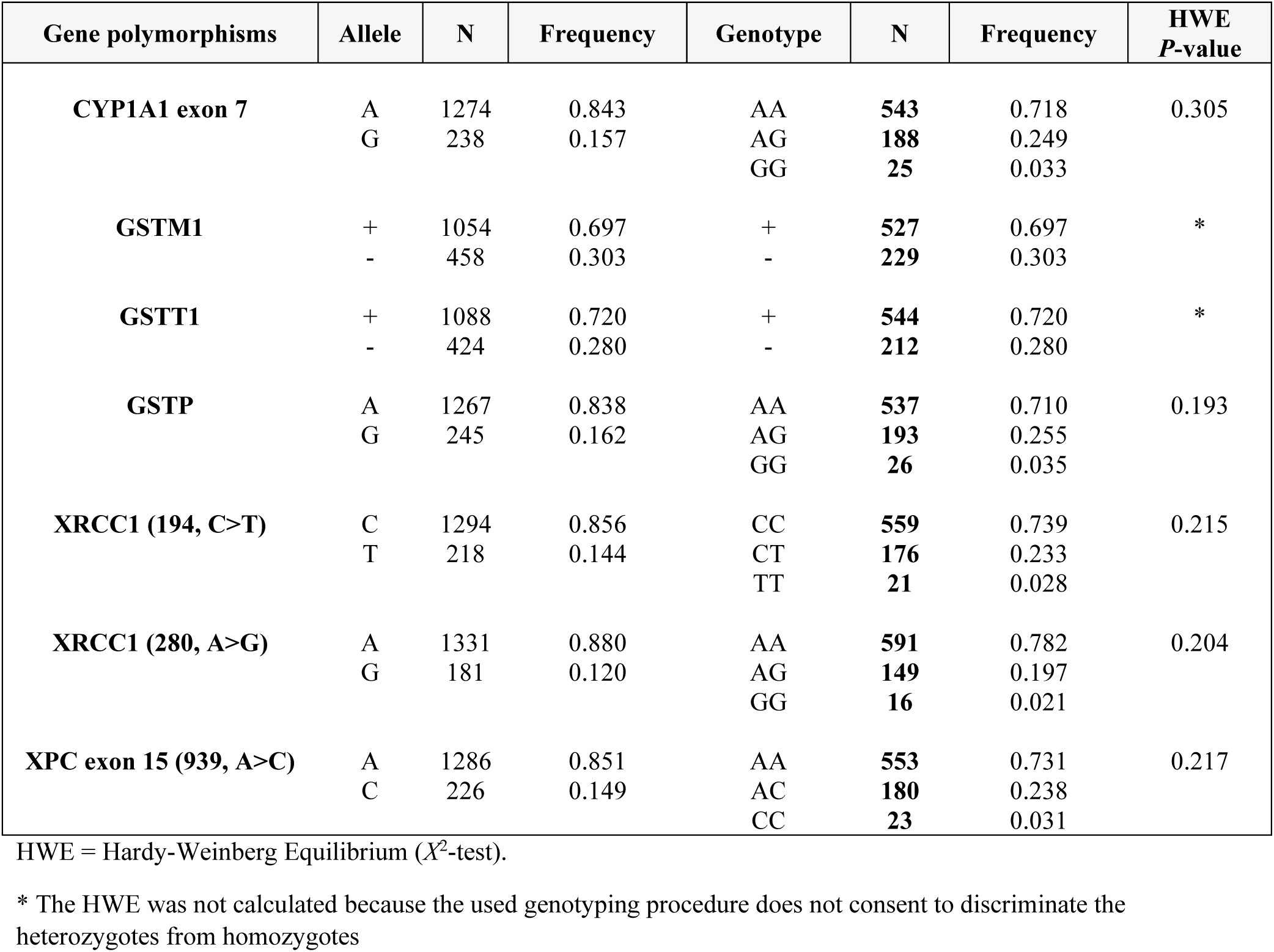
– Allele and Genotype Frequencies of seven gene polymorphisms in an Italian sample (n = 756)

| Gene polymorphisms | Allele | N | Frequency | Genotype | N | Frequency | HWE<br>*P*-value |
| --- | --- | --- | --- | --- | --- | --- | --- |
| **CYP1A1 exon 7** | A | 1274 | 0.843 | AA | **543** | 0.718 | 0.305 |
|  | G | 238 | 0.157 | AG | **188** | 0.249 |  |
|  |  |  |  | GG | **25** | 0.033 |  |
| **GSTM1** | + | 1054 | 0.697 | + | **527** | 0.697 | * |
|  | - | 458 | 0.303 | - | **229** | 0.303 |  |
| **GSTT1** | + | 1088 | 0.720 | + | **544** | 0.720 | * |
|  | - | 424 | 0.280 | - | **212** | 0.280 |  |
| **GSTP** | A | 1267 | 0.838 | AA | **537** | 0.710 | 0.193 |
|  | G | 245 | 0.162 | AG | **193** | 0.255 |  |
|  |  |  |  | GG | **26** | 0.035 |  |
| **XRCC1 (194, C>T)** | C | 1294 | 0.856 | CC | **559** | 0.739 | 0.215 |
|  | T | 218 | 0.144 | CT | **176** | 0.233 |  |
|  |  |  |  | TT | **21** | 0.028 |  |
| **XRCC1 (280, A>G)** | A | 1331 | 0.880 | AA | **591** | 0.782 | 0.204 |
|  | G | 181 | 0.120 | AG | **149** | 0.197 |  |
|  |  |  |  | GG | **16** | 0.021 |  |
| **XPC exon 15 (939, A>C)** | A | 1286 | 0.851 | AA | **553** | 0.731 | 0.217 |
|  | C | 226 | 0.149 | AC | **180** | 0.238 |  |
|  |  |  |  | CC | **23** | 0.031 |  |

HWE = Hardy-Weinberg Equilibrium ( $X^2$ -test).
\* The HWE was not calculated because the used genotyping procedure does not consent to discriminate the heterozygotes from homozygotes

The frequencies of the metabolic and DNA-repair gene polymorphisms among age groups are reported in Table 4. We observed a general decreasing trend in the frequency of *GSTT1* null allele across all age groups. In particular, we observed a significant decrease in the frequency of the null allele in the oldest group with respect to the youngest one (*P =* 3.72 × 10^−2^) and with respect to 51-85 age group (*P =* 8 × 10^−4^). This last age group showed a significant higher frequency of null allele with respect to 18-50 age group (*P =* 2 × 10^−2^).

**Table 4.**
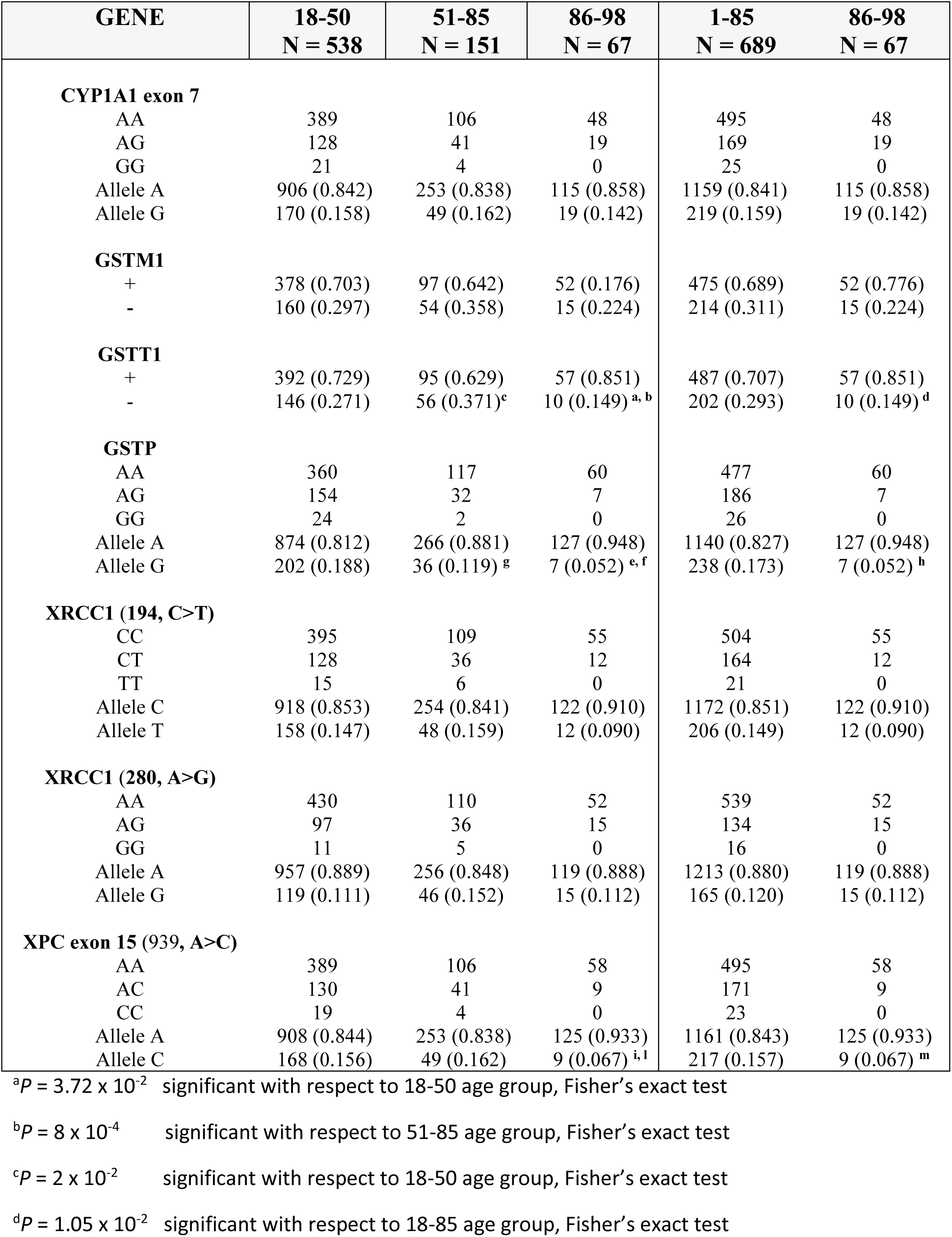
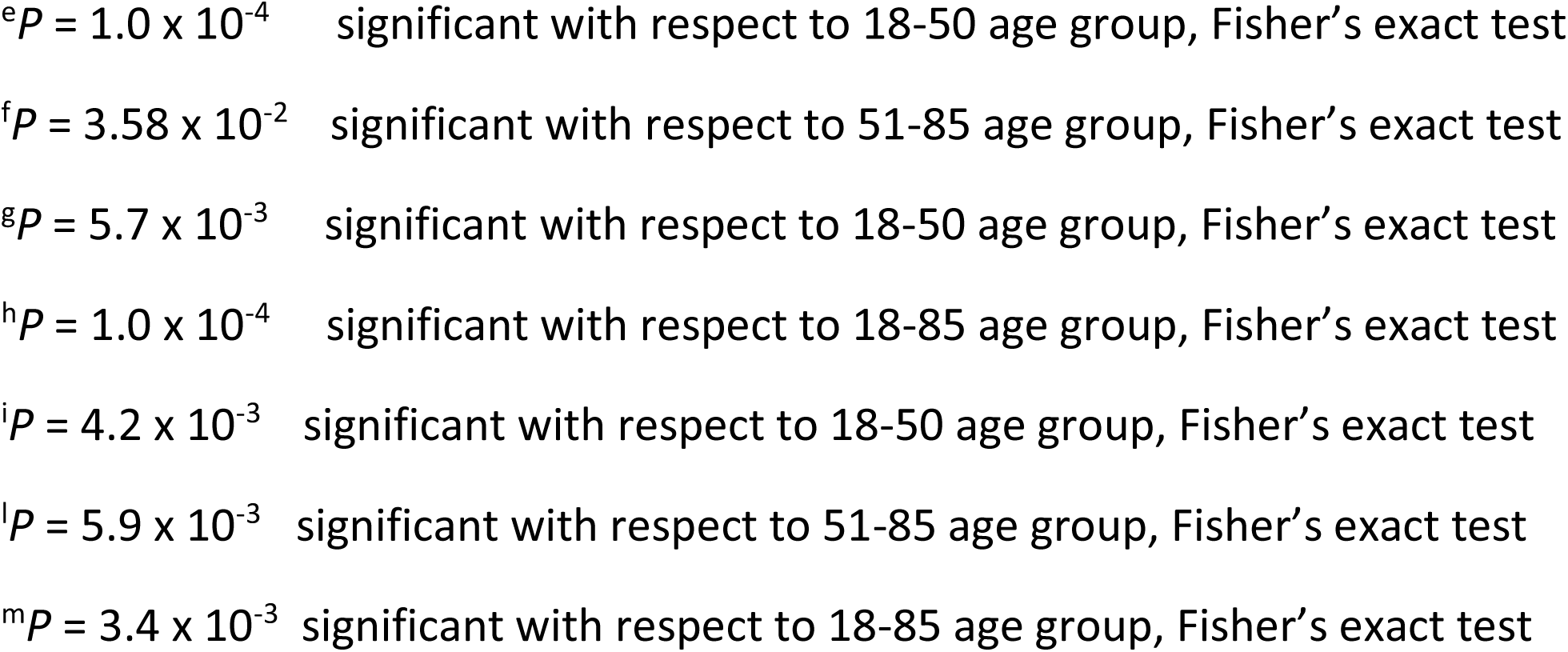
Frequencies of the studied metabolic gene polymorphisms among different age groups (n = 756). Only significant results were reported.

| GENE | 18-50<br>N = 538 | 51-85<br>N = 151 | 86-98<br>N = 67 | 1-85<br>N = 689 | 86-98<br>N = 67 |
| --- | --- | --- | --- | --- | --- |
| **CYP1A1 exon 7** |  |  |  |  |  |
| AA | 389 | 106 | 48 | 495 | 48 |
| AG | 128 | 41 | 19 | 169 | 19 |
| GG | 21 | 4 | 0 | 25 | 0 |
| Allele A | 906 (0.842) | 253 (0.838) | 115 (0.858) | 1159 (0.841) | 115 (0.858) |
| Allele G | 170 (0.158) | 49 (0.162) | 19 (0.142) | 219 (0.159) | 19 (0.142) |
| **GSTM1** |  |  |  |  |  |
| + | 378 (0.703) | 97 (0.642) | 52 (0.176) | 475 (0.689) | 52 (0.776) |
| - | 160 (0.297) | 54 (0.358) | 15 (0.224) | 214 (0.311) | 15 (0.224) |
| **GSTT1** |  |  |  |  |  |
| + | 392 (0.729) | 95 (0.629) | 57 (0.851) | 487 (0.707) | 57 (0.851) |
| - | 146 (0.271) | 56 (0.371)<sup>c</sup> | 10 (0.149)<sup>a, b</sup> | 202 (0.293) | 10 (0.149)<sup>d</sup> |
| **GSTP** |  |  |  |  |  |
| AA | 360 | 117 | 60 | 477 | 60 |
| AG | 154 | 32 | 7 | 186 | 7 |
| GG | 24 | 2 | 0 | 26 | 0 |
| Allele A | 874 (0.812) | 266 (0.881) | 127 (0.948) | 1140 (0.827) | 127 (0.948) |
| Allele G | 202 (0.188) | 36 (0.119)<sup>g</sup> | 7 (0.052)<sup>e, f</sup> | 238 (0.173) | 7 (0.052)<sup>h</sup> |
| **XRCC1 (194, C>T)** |  |  |  |  |  |
| CC | 395 | 109 | 55 | 504 | 55 |
| CT | 128 | 36 | 12 | 164 | 12 |
| TT | 15 | 6 | 0 | 21 | 0 |
| Allele C | 918 (0.853) | 254 (0.841) | 122 (0.910) | 1172 (0.851) | 122 (0.910) |
| Allele T | 158 (0.147) | 48 (0.159) | 12 (0.090) | 206 (0.149) | 12 (0.090) |
| **XRCC1 (280, A>G)** |  |  |  |  |  |
| AA | 430 | 110 | 52 | 539 | 52 |
| AG | 97 | 36 | 15 | 134 | 15 |
| GG | 11 | 5 | 0 | 16 | 0 |
| Allele A | 957 (0.889) | 256 (0.848) | 119 (0.888) | 1213 (0.880) | 119 (0.888) |
| Allele G | 119 (0.111) | 46 (0.152) | 15 (0.112) | 165 (0.120) | 15 (0.112) |
| **XPC exon 15 (939, A>C)** |  |  |  |  |  |
| AA | 389 | 106 | 58 | 495 | 58 |
| AC | 130 | 41 | 9 | 171 | 9 |
| CC | 19 | 4 | 0 | 23 | 0 |
| Allele A | 908 (0.844) | 253 (0.838) | 125 (0.933) | 1161 (0.843) | 125 (0.933) |
| Allele C | 168 (0.156) | 49 (0.162) | 9 (0.067)<sup>i, l</sup> | 217 (0.157) | 9 (0.067)<sup>m</sup> |

<sup>a</sup>P = 3.72 x 10<sup>-2</sup> significant with respect to 18-50 age group, Fisher's exact test
<sup>b</sup>P = 8 x 10<sup>-4</sup> significant with respect to 51-85 age group, Fisher's exact test
<sup>c</sup>P = 2 x 10<sup>-2</sup> significant with respect to 18-50 age group, Fisher's exact test
<sup>d</sup>P = 1.05 x 10<sup>-2</sup> significant with respect to 18-85 age group, Fisher's exact test
<sup>e</sup> $P = 1.0 \times 10^{-4}$ significant with respect to 18-50 age group, Fisher's exact test
<sup>f</sup> $P = 3.58 \times 10^{-2}$ significant with respect to 51-85 age group, Fisher's exact test
<sup>g</sup> $P = 5.7 \times 10^{-3}$ significant with respect to 18-50 age group, Fisher's exact test
<sup>h</sup> $P = 1.0 \times 10^{-4}$ significant with respect to 18-85 age group, Fisher's exact test
<sup>i</sup> $P = 4.2 \times 10^{-3}$ significant with respect to 18-50 age group, Fisher's exact test
<sup>l</sup> $P = 5.9 \times 10^{-3}$ significant with respect to 51-85 age group, Fisher's exact test
<sup>m</sup> $P = 3.4 \times 10^{-3}$ significant with respect to 18-85 age group, Fisher's exact test

Similarly, for *GSTP* gene polymorphism, the oldest age group showed a significant lower value of the minor allele with respect to 18-50 (*P* = 1 × 10^−4^) and 51-85 (*P* = 1 × 10^−4^) age groups, as well as this last age group showed a significant lower frequency of the G-allele with respect to 18-50 age group (*P =* 5.7 × 10^−3^). For XPC exon 15 A939C gene polymorphisms, we observed a significant decrease of the C-allele in the oldest age group with respect to 18-50 (*P =* 4.2 × 10^−3^) and 51-85 (*P =* 5.9 × 10^−3^) age groups.

Also subdividing the sample in two age groups, 18-85 and 86-98, we obtained the same results, with the oldest age group showing a significant decrease of the *GSTT1* null (*P* = 1.05 × 10^−2^), *GSTP* G (*P* = 1 × 10^−4^) and XPC C (*P* = 3.4 × 10^−3^) allele frequencies with respect to 18-85 age group.

Finally, the multivariate general linear model confirmed this significantly decreasing trend with age for *GSTT* null (*P* = 3.2 × 10^−2^), *GSTP* G (*P* = 8 × 10^−3^) and *XPC* C (*P* = 3.4 × 10^−2^) alleles.

## 4. Discussion

Longevity is due to a complex interaction of genetic and environmental factors, but a strong genetic component appears to have an impact on survival to extreme ages. Indeed, heritability estimates of longevity suggest that about a third of the phenotypic variation associated with the trait is attributable to genetic factors, and the rest is influenced by epigenetic and environmental factors (Govindaraju et al., 2015).

In order to identify longevity genes in humans, different strategies can be adopted. In the present paper we performed an association study, conducted on an Italian sample and analyzing seven drug and DNA-repair gene polymorphisms. We found a negative association of *GSTT1 null, GSTP1 G* and *EPC* exon 15 C alleles with human longevity.

A possible explanation of our findings is that individual genetic susceptibility play an increasingly important role in determining the levels of genomic damage, as well as the studied gene polymorphisms were found to be associated with different type of cancers. In particular, GSTT1 and GSTP1 polymorphic enzymes, involved in the processing of reactive oxygen, lipid peroxidation products and some key metabolites of toxicants, may change metabolic process efficiency, which contribute to the individual disease susceptibility (Bolt & Thier, 2006). Many studies have shown that variants of *GSTs* are significantly associated with disease risk, including different types of cancers, cardiovascular and allergic diseases (Di Pietro et al., 2010; Eslami and Sahebkar, 2014; Dar et al., 2017). Genetic variant in *GSTT1* consists of a deletion of the whole gene, resulting in the lack of the active enzyme, making the phenotypic variation in glutathione related detoxification negative (*GSTT1 null*) (Di Pietro et al., 2010). Consequently, the lowest frequency of *GSTT1 null* genotypes found in subjects aged between 86 and 98 years seems to suggest a selective disadvantage of possible detrimental genotypes in this age group. These data are in line with results published by Santovito et al. (2008), who found a low frequency of the *GSTT1 null* genotype in subjects aged between 80 and 100.

Like *GSTT1*, also *GSTP1* has a corresponding rationale for an association with age. GSTP1 enzyme performs functions such as metabolism of halogenated compounds, of low molecular weight molecules and reactive epoxides. On codon 105, which comprises part of the active site of *GSTP1*, the transition from adenine to guanine can occur (Chielle et al., 2016). When this exchange takes place, substrate-specific catalytic activity becomes substantially less effective in its activity, resulting in the development of chronic diseases such as cardiovascular diseases and some types malignant neoplasms (Khabaz, 2014; Chielle et al., 2016).

We found no differences between age groups in the frequency of *CYP1A1 exon 7* genotype. These data seem to be concordant with other studies, such as the one conducted by Taioli et al. (2001), that did not find any difference in the frequency of *CYP1A1* in centenarians, as compared to younger control subjects. However, published data evaluating the possible association between phase I enzyme polymorphisms and ageing reported conflicting results (Wuathier et al., 2007; Seripa et al., 2010).

Finally, distinct mechanisms have evolved to repair different types of DNA damage and to maintain genomic integrity. Several researches have demonstrated that subjects showing compromised repair capacity have increased mutation rates and genomic instability, and, consequently, an increased risk of cancer and a reduced life-expectancy (Cornetta et al 2006). In general, healthy persons differ in intrinsic capacity in repairing DNA damage, and this variation may results from polymorphisms in genes involved in different repair pathways. Several epidemiologic studies have highlighted a role for some SNPs in NER and BER genes as risk factors for cancer (Fontana et al., 2008). In this scenario, we hypothesized that polymorphisms in *XPC* gene, being important in the sensitivity against the development of different types of cancer (Bahceci et al., 2015), may reflect a reduced life-expectancy in carrier subjects and may explain the significant low frequency of *XPC* exon 15 C-allele observed in the present study among subjects belonging to oldest age group. Conversely, there is no consensus regarding the possible association between *XRCC1* Arg194Trp and *XRCC1* Arg280His with different types of cancer (Moghaddam et al., 2016; Dylawerska et al., 2017), data that may explain the lack of association of these gene polymorphisms with longevity, as observed in our study.

## Conclusions

With this study we highlighted a role for *GSTT1, GSTP* and *XPC* gene polymorphisms in longevity, also evidencing the utility of a comparative analysis of longevity between individuals with normal life span and elderly subjects. Although these results cannot be generalized, due to the different genetic background of the worldwide distributed populations, as well as to the different environment in which they live, data about frequencies of cytokine gene polymorphisms in different age classes, could be useful to better clarify the role of these gene polymorphisms in longevity-related processes.

However, we would like to emphasize some limitations typical of association studies. For example, populations could be heterogeneous in terms of genetic ancestry and, may comprise two or more groups with distinct genetic ancestry (Hellwege et al., 2017; Liu et al., 2013). This population stratification is principally due to non-random mating processes following to geographic isolation of subpopulations with low rates of migration and gene flow (Hellwege et al., 2017) and it can give rise to a spurious association of one or more alleles with longevity.

Finally, the subdivision of the sample into age groups poses the problem of possible early life differences in mortality in some cohorts. As consequence, elderly subjects could have different infant mortality rates than younger subjects. This situation could lead to a false association of some alleles with longevity while, in reality, the association could be simply due to a differential survival in the early stages of life.

